# Effects of L-Dopa, SKF-38393 and Quinpirole on exploratory, anxiety- and depressive-like behaviors in pubertal female and male mice

**DOI:** 10.1101/2021.02.26.433029

**Authors:** Muiara Aparecida Moraes, Laila Blanc Árabe, Bruna Lopes Resende, Beatriz Campos Codo, Ana Luiza de Araújo Lima Reis, Bruno Rezende de Souza

**Affiliations:** Laboratório de Neurodesenvolvimento e Evolução - Department of Physiology and Biophysics, Federal University of Minas Gerais, Belo Horizonte, MG, 31270-901, Brazil

**Keywords:** Dopamine, D1, D2, sexual dimorphism, ontogeny, adolescence

## Abstract

Adolescence is a phase of substantial changes in the brain, characterized by maturational remodeling of many systems. This remodeling allows functional plasticity to adapt in a changing environment but turns this period into a neurodevelopmental vulnerable window. The dopaminergic system is under morphological and physiological changes during this phase. The disruption of its balance can lead to molecular variation and abnormal behavior - representing a risk factor for neuropsychiatric disorders. In the present study, we investigated if changes in the dopaminergic tone alter mice behavior in a receptor and sex-specific manner, specifically in the beginning of puberty period. We administered L-Dopa, SKF-38393 (D1 dopamine receptor agonist) and Quinpirole (D2 dopamine receptor agonist) and tested male and female mice motor, anxiety- and depressive-like behavior. While females displayed an impaired exploratory drive, males presented an intense depressive-like response. Our results provide insights into the function of dopaminergic development in adolescent behavior and highlight the importance of studies in this time window with male and female subjects.

## Introduction

Adolescence is a phase of extensive changes in the brain, marked by functional, chemical, and neuroanatomical reorganization (Keshavan et al., 2014; Shaw et al., 2008). In particular, the dopaminergic system undergoes substantial remodeling (Wahlstrom et al. 2010). Although dopamine (DA) innervation is present since earlier in development, several studies demonstrated that mesocortical projections increase during adolescence, in both rodents and primates (Hoops et al., 2018; Hoops and Flores, 2017; Kalsbeek et al., 1988; Naneix et al., 2012; Reynolds et al., 2019; Rosenberg and Lewis, 1994; Willing et al., 2017). This protracted development of mesocortical DA innervation may have an essential role during the maturation of behavior, and many molecular patterns are still maturing in adolescent DAergic signaling. For example, even though adolescent animals can make value-based decisions, they are not able to adapt actions according to contingency changes (Naneix et al., 2012), similarly to adult animals with lesions of dopaminergic innervation in the prefrontal cortex (Naneix et al., 2009). Baseline tissue levels of DA also differ during adolescence in a region-dependent manner (Corongiu et al., 2019), as well as dopamine transporter (DAT) functionality and DA release following electrical stimulation (Walker and Kuhn, 2008).

Brain remodeling during adolescence allows adaptation to the environmental context but also turns this window into a vulnerable period to risk factors for psychiatric disorders. Adolescence has been described as a period of vulnerability to drug abuse (Corongiu et al., 2019; Cservenka et al., 2014; Squeglia & Cservenka, 2017), and several psychiatric disorders emerge at this time point (Kessler et al., 2007; Merikangas et al., 2009; Breslau et al., 2017). Major depression disorder diagnoses peak in early adolescence, with higher rates of suicide attempts during this period; and women are more diagnosed in all developmental periods (Breslau et al., 2017; Hankin et al., 1998; Rohde et al., 2013). There is evidence of brain functional and anatomical differences in schizophrenia patients even before the emergence of symptoms (Johnstone et al., 2005; Murray & Lewis, 1987; Thermenos et al., 2013), which appear earlier and with more negative traits in men than in women (Abel et al., 2010). Interestingly, the DAergic system plays a role in the pathophysiology of these disorders. For example, major depression is associated with an increase of DA receptors and reduced DA transmission (Dunlop & Nemeroff, 2007; Grace, 2016). Exploring the neurophysiology of early adolescence, before the onset of psychiatric disorders, is crucial to understand their etiology.

In the mammalian brain, DA signaling occurs via five metabotropic receptors that divide into two major groups: D1 and D2 family, and when activated those receptors lead to opposing changes in intracellular cascades (Beaulieu & Gainetdinov, 2011). The distribution of DA receptors in the brain is heterogeneous - with distinct proportions and concentrations according to the brain areas (Andersen and Teicher, 2000; Hurd et al., 2001; Tarazi & Baldessarini, 2000). The ratio of D1: D2 receptors also distinguishes according to region, age, or sex (Cullity et al., 2019), and this balance is suggested to influence behavioral responses. For example, cue-associated learning extinction behavior is partially impaired in adolescent rats, and the administration of quinpirole, a D2 agonist, in the infralimbic cortex, seems to enhance long term memory extinction in those animals (Zbukvic et al., 2017; Zbukvic and Hyun Kim, 2018). Interestingly, photostimulation in D1, but not D2 expressing neurons in the medial prefrontal cortex, led to antidepressant and anxiolytic effects (Hare et al., 2019). On the other hand, photostimulation in D1 and D2 expressing neurons in the nucleus accumbens led to enhanced motivation in mice (Soares-Cunha et al., 2016). These findings suggest a region- and behavior-specific effect. Specifically, during adolescence, D1 receptors are pivotal for social behavior shaping (Kopec et al., 2018). Thus, DA signaling is a determinant factor for many behavioral outcomes, and interventions in this system can lead to different patterns of behavior (Areal and Blakely, 2020).

The DAergic system also differs between sex. Receptor rates fluctuate in a sex and region-specific manner throughout maturation (Andersen et al., 1997; Andersen & Teicher, 2000; Orendain-Jaime et al., 2016), and females have a higher D1: D2 ratio compared to males (Cullity et al., 2019). Kopec et al. (2018) demonstrated that mesolimbic maturation, specifically in the nucleus accumbens, occurs differently in male and female mice (Kopec et al., 2018). Sannino et al. (2017) suggested adolescence as a sex-related watershed for behavioral and molecular abnormalities caused by variations in an enzyme that degrade DA (Sannino et al., 2017). Furthermore, early life disruptions alter DAergic function differently in males and females (Dwyer & Leslie, 2016; Kawakami et al., 2013; Stewart et al., 1991). A transient postnatal increase of DA leads to different behavior changes in male and female pubertal and adult mice (de Matos et al., 2018, 2020). Thus, adding female subjects to studies on puberty/adolescent time-window is relevant to understand better the differences seen in some psychopathologies.

Changes in the DAergic system during vulnerable developmental windows can lead to molecular variations and abnormal behavior. Early adolescence, or the beginning of puberty, is a vulnerable period, marked by a refinement of circuits, including the DAergic pathways, and the expression of DA receptors during this stage diverges in a sexually different manner. Taking all these into consideration, we hypothesized that changes in the prepubertal mice DAergic tone alter brain function and behavior in a receptor and sex-specific manner. This study is the first to our knowledge to investigate the consequences of the pharmacological activation of D1 and D2 receptors during early adolescence in anxiety- and depressive-like behaviors of female and male mice.

## Materials

### Animals

We purchased male and female Swiss mice (Tac:UFMG: SW / 8-12 weeks of age) from the Animal Facility of the Federal University of Minas Gerais (CEBIO, UFMG, Brazil) for mating. Mice were raised in polypropylene cages (27cm x 17cm x 12cm), provided with wood shavings, housed three to five per cage, maintained in a climate-controlled room with 12h-12h light/dark cycle (lights on from 7 am to 7 pm), temperature at 22+-2 °C and 40-70% relative humidity. Free access to food (Nuvilab CR1 - Nuvital) and water was provided throughout the study.

We conducted all protocols during the light phase of the cycle. We performed the behavioral protocols in pubertal (28-32 days old) male and female Swiss mice offspring (Nelson et al., 1990). The Animal Use Ethics Committee of the Federal University of Minas Gerais (CEUA 39/2015) approved all procedures, and the experiments were carried out following NIH guidelines for the use and care of animals (Liu et al., 2011).

### Experimental design and drug administration

We stimulated mating by placing two females with one male. After five days, the male was removed and two females were housed per cage. On the 19th day of gestation, each female was housed individually. We then inspected the cages daily for the presence of pups. The day of birth was designated PD1, and litter-size was standardized in 8 to 9 pups. At PD21, the animals were weaned and housed by sex. From this day on, an experimenter habituated the animals, handling them in the same manner to the day of behavior tests. At P30, the animals were randomly assigned to their treatment groups. They received intraperitoneal injections of saline for control, the dopamine precursor L-Dopa + benserazide (10/5 and 50/25 mg/kg; Sigma Aldrich D1507 and B7283, respectively), the D2 agonist quinpirole (0,5 mg/Kg Sigma Aldrich Q111) or the D1 agonist SKF 38393 (5mg/Kg Sigma Aldrich D047); the volume of injection was 11,67ml/Kg (Arnauld et al., 1993; Beninger & Ranaldi, 1992; de Matos et al., 2018; Eilam & Szechtman, 1989; Luque-Rojas et al., 2013). The animals were submitted to an open field test and elevated plus maze 30 min after injections or to forced swimming test 50 min after injections.

### Behavior tests

#### Open Field Test

We used the Open Field Test to measure exploratory behavior and locomotor activity in rodents. Mice were placed in an automatic open field apparatus (LE 8811 IR Motor Activity Monitors PANLAB, Harvard Apparatus; Spain), with an acrylic box measuring 23×23×35cm. Animals were free to explore the field for 20 min. We recorded the total distance travelled, and the percentage of time spent in the center of the open field (10 x 10 cm) was used as an anxiety measure (Prut & Belzung, 2003; Simon et al., 1994; Walsh & Cummins, 1976). Rearing and increased locomotion were considered a stimulant effect (Prut and Belzung, 2003). The open-field arena was cleaned with 70% ethanol between each animal.

#### Elevated Plus Maze

We used the Elevated Plus Maze to identify anxiety-related behavior in rodents (Walf & Frye, 2007). The apparatus consists of two open arms (30cm×6cmx16cm), and two closed arms (30cm×6cm×16cm) crossed in the middle perpendicularly to each other and elevated 30cm above the floor. Mice were placed in the central area and allowed to move freely for 5 minutes, due to a highly responsive behavior over this time (Montgomery, 1955). We used the percentage of entries into the open arms and the time spent in the open arms as indices of anxiety (Pellow et al., 1985). We also analyzed ethological parameters such as rearings, stretchings and head-dippings in order to explore the complexity of mouse behavior (Espejo, 1997). We counted vertical exploration as rearings, projection of mice’s head over the edge of the open arms, towards the floor, as head-dippings, and the stretching of the forepaws without moving the hind paws as stretchings (Rodgers et al., 1997; Rodgers & Johnson, 1995).

#### Forced Swimming Test

We used the Forced Swimming Test to evaluate depressive-like behavior in rodents, and it was performed as reviewed by Can and colleagues in 2012, with minor modifications (Can et al., 2011; Porsolt et al., 1977). Mice were individually transferred to a vertical glass cylinder (17 cm inner diameter × 27 cm height) filled with water, and were forced to swim for 6 minutes. The water’s temperature was monitored throughout the test and maintained between 25 and 27 C°. We recorded and analyzed the entire test. Immobility was quantified when mice were merely floating, making only movements necessary to keep the head above the water. We quantified climbing as vertical movements against the cylinder wall. Activity time was defined subtracting immobility time from the total time. Swimming time was defined subtracting the climbing time from the activity time. The immobility rate was defined as immobility time per number of immobility episodes.

#### Statistical Analysis

Parametric data were analyzed by One Way ANOVA followed by Tukey’s posthoc test. Non-parametric data were analyzed by Kruskal-Wallis One Way Analysis of Variance on Ranks, followed by Dunn’s Method posthoc test. Parametric data are presented as mean and standard deviation (SD). Non-parametric data are presented as median and interquartile (Weissgerber et al., 2015). Significance was set at p < 0.05 for all analyses.

To evaluate whether there was a difference in results between sexes, we normalized female and male data by their control (Saline). We then performed Two-Way ANOVA followed by Tukey’s posthoc test. Data are presented as mean and standard deviation (SD).

## Results

### Acute interference in dopaminergic signaling leads to sex-specific and exploratory-related locomotor changes

Alterations in the dopaminergic system are related to parallel changes in locomotor activity (Dunnett, 2005). In our previous work, we showed that increasing dopamine levels during the first days of life affect the locomotor activity of pubertal and adult female mice (de Matos et al., 2020; 2018). In this study, we investigate if an acute injection of L-Dopa or dopamine receptor D1 and D2 agonists during the prepubertal period can change motor behavior. We observed no alterations in the locomotor activity of pubertal males (Figure 1A and C); and we found a significant decrease in the total distance travelled in L-Dopa 50mg/Kg injected females. The L-Dopa 50mg/Kg injected females moved around approximately 31% less than the saline group (KW H=13.526, 4 d.f., Dunn’s Method, p<0.05) (Figure 1A). When we analyzed every 5 minutes, we noticed that L-Dopa 50mg/kg females only walk less during the first 5 minutes (F(16,295)= 1,255, Two Way ANOVA, Tukey, p= 0.002, figure 1B and KW H=14.692, 4 d.f., Dunn’s Method, p<0.05, figure 1B’), and not during the whole test. Also, L-Dopa 10mg/Kg females spent 31% less time in the center when compared to the control group (KW H=10.691, 4 d.f., Dunn’s Method, p<0.05) (Figure 1D). There was a decrease of 33% in the number of rearings in the L-Dopa 10mg/Kg group (KW H=48.478, 4 d.f., Dunn’s Method, p<0.05) (Figure 1E), as well as a decrease of 53% in the L-Dopa 50m/Kg group (p<0.05) (Figure 1E). Additionally, Quinpirole injected females decreased the number of rearings when compared to saline females (p<0.05) (Figure 1E).

**Fig. 1.**
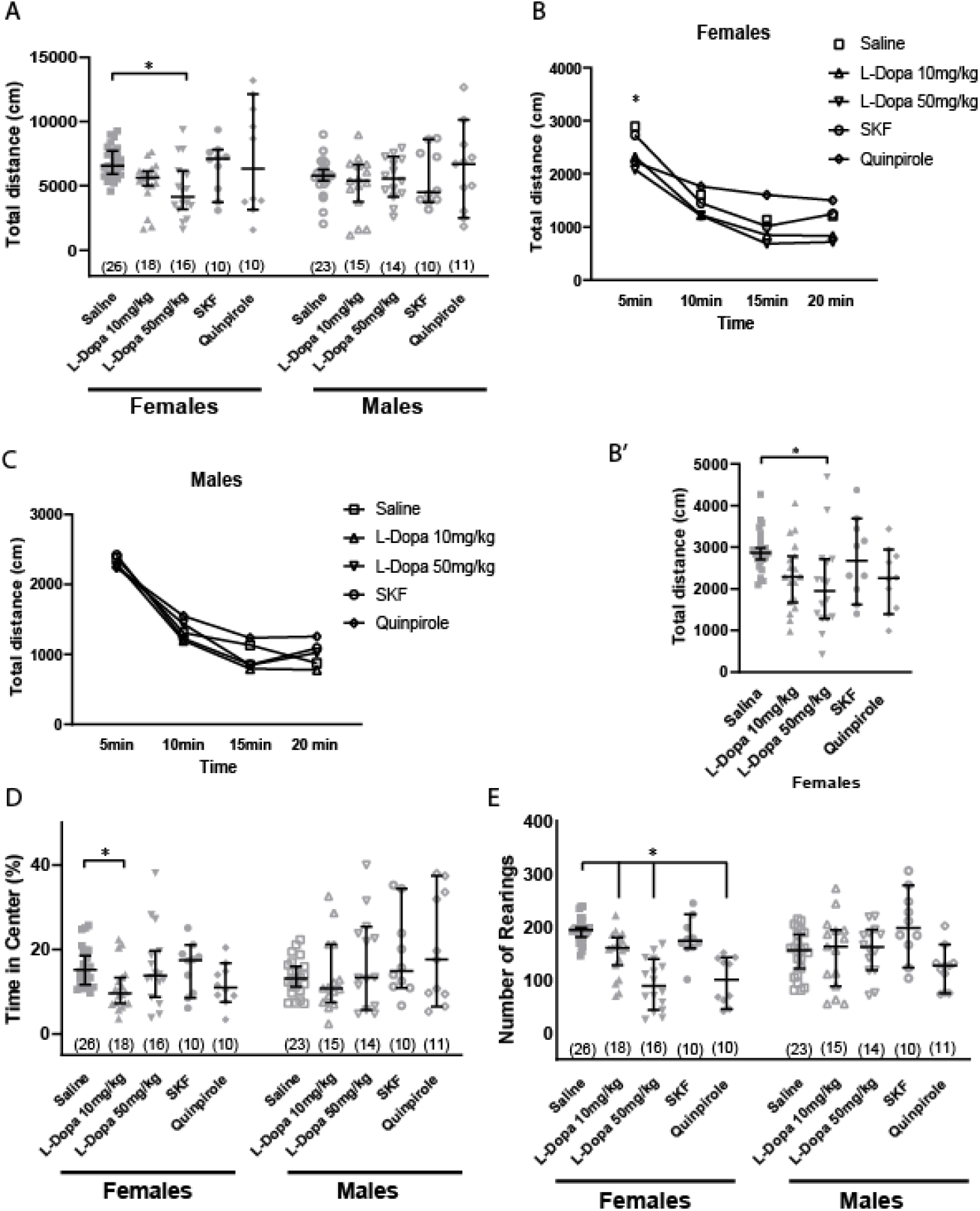
Effects of an acute increase of dopaminergic signaling during adolescence in exploratory-driven locomotion. Motor behavior of prepubertal Swiss mice (PD30) after injection of Saline, L-Dopa 10mg/kg, L-Dopa 50mg/kg, SKF-38393 or Quinpirole tested in the Open Field Test. (A) We observed alterations in the total distance travelled only in L-Dopa 50mg/kg females. (B) To analyze the differences between the groups over time, we used the Two Way ANOVA and Tukey Test. We found that L-Dopa 50mg/kg females showed a reduced locomotion only in the first interval, (B’) as confirmed by a One Way ANOVA followed by Tukey Test. (C) Males from all experimental groups showed a similar behavior during the whole test. (D) L-Dopa 10mg/kg females spent less time in the center of the Open Field compared to control. (E) L-Dopa 10mg/kg, L-Dopa 50mg/kg and Quinpirole females had a reduction in the number of rearings compared to Saline. The number of animals is between parentheses. Nonparametric data are represented as median with interquartile range. Parametric data are represented as mean ± SD. For parametric data we used One Way ANOVA test and Tukey Test. For nonparametric data we used Kruskal-Wallis One Way ANOVA on Ranks and Dunn’s test. *p < 0.05.

Then we tested whether the effects in the locomotor activity arising from the acute L-Dopa and agonists administration were sex-specific. We observed that the difference between sexes was statistically significant in the total distance walked within the L-Dopa 50mg/kg group (F(4,143)= 0.876, Two Way ANOVA, Tukey, p=0.022) (Figure 2A). Quinpirole injected males and females spent a different amount of time in the center of the box (F(4,143)= 1.844, Two Way ANOVA, Tukey, p=0.002) (Figure 2B). As for the number of rearings, L-Dopa 10mg/kg (F(4,143)= 5.377, Two Way ANOVA, Tukey, p=0.017), L-Dopa 50mg/kg (p<0.001), SKF (p<0.001) and Quinpirole (p=0.013) groups showed differences in a sex-specific manner (Figure 2C).

**Fig. 2.**
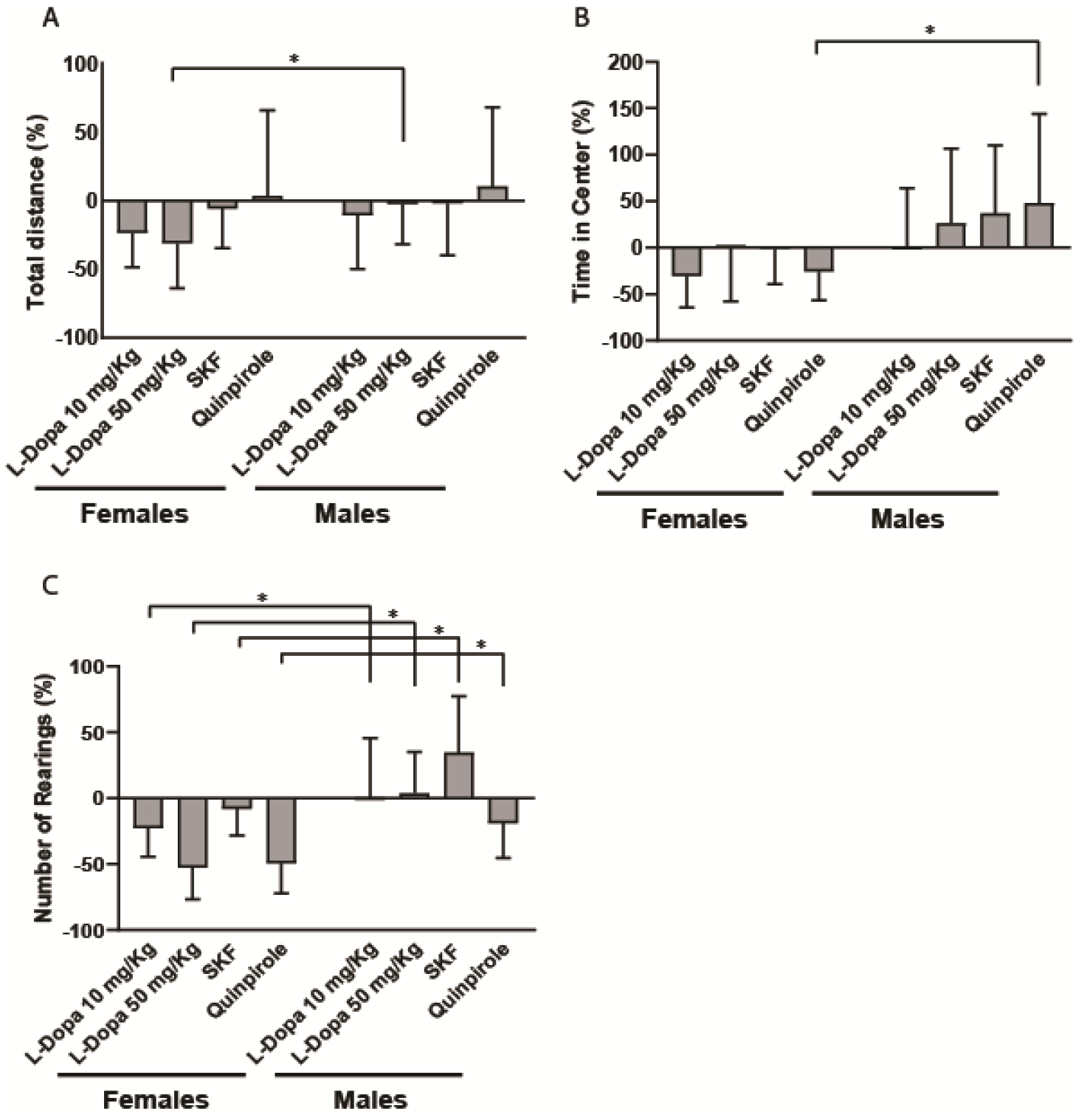
Locomotor sex-specific effects after an acute increase of dopaminergic signaling during pubertal phase. Motor behavioral differences between male and female prepubertal mice injected with Saline, L-Dopa 10mg/kg, L-Dopa 50mg/kg, SKF-38393 or Quinpirole. (A) The total distance travelled by L-Dopa 50mg/kg males was higher than the travelled by females. (B) Quinpirole males spent more time in the center of the Open Field compared to females of the same group. (C) The number of rearing was higher in males from all experimental groups compared to females from the respective group. We used the Two Way ANOVA and Tukey Test, data are represented as mean ± SD *p < 0.05.

### Increased D2 receptor stimulation led to a decrease of exploratory behavior in mice

Considering the critical role of dopaminergic signaling as a modulator for anxiety-like behaviors (Calhoon and Tye, 2015; de la Mora et al., 2010; Pezze and Feldon, 2004), we used the Elevated Plus Maze to test whether the administration of L-Dopa or dopamine receptor D1 and D2 agonists alter anxiety-like behavior at the beginning of the pubertal phase in females and males. We observed no alterations in the percentage of entries or time in the open arms, neither in females or males (Figure 3A and B). There were also no changes in time in the center or in the open arms (Figure C). L-Dopa 50mg/Kg females showed a decrease of 42% in the number of rearings (F(4,78)=19.591, One Way Anova, Tukey, p<0.001), and 68% for the Quinpirole injected females (p<0.001) when compared to the saline group (Figure 3D). We also observed a reduction of 33% in the number of dippings in females administered with L-Dopa 50mg/Kg (KW H=27.334, 4 d.f., Dunn’s Method, p<0.05), a 40% decrease in the SKF group (p<0.05) and a 59% decrease in the Quinpirole group (p<0.05), when compared to saline females (Figure 3E). Furthermore, we found a reduction of 33% in the number of stretchings in L-Dopa 10mg/Kg (KW H=34.473, 4 d.f., Dunn’s Method, p<0.05) and a decrease of 47% in L-Dopa 50mg/Kg females (p<0.05) compared to the control group (Figure 3F).

**Fig. 3.**
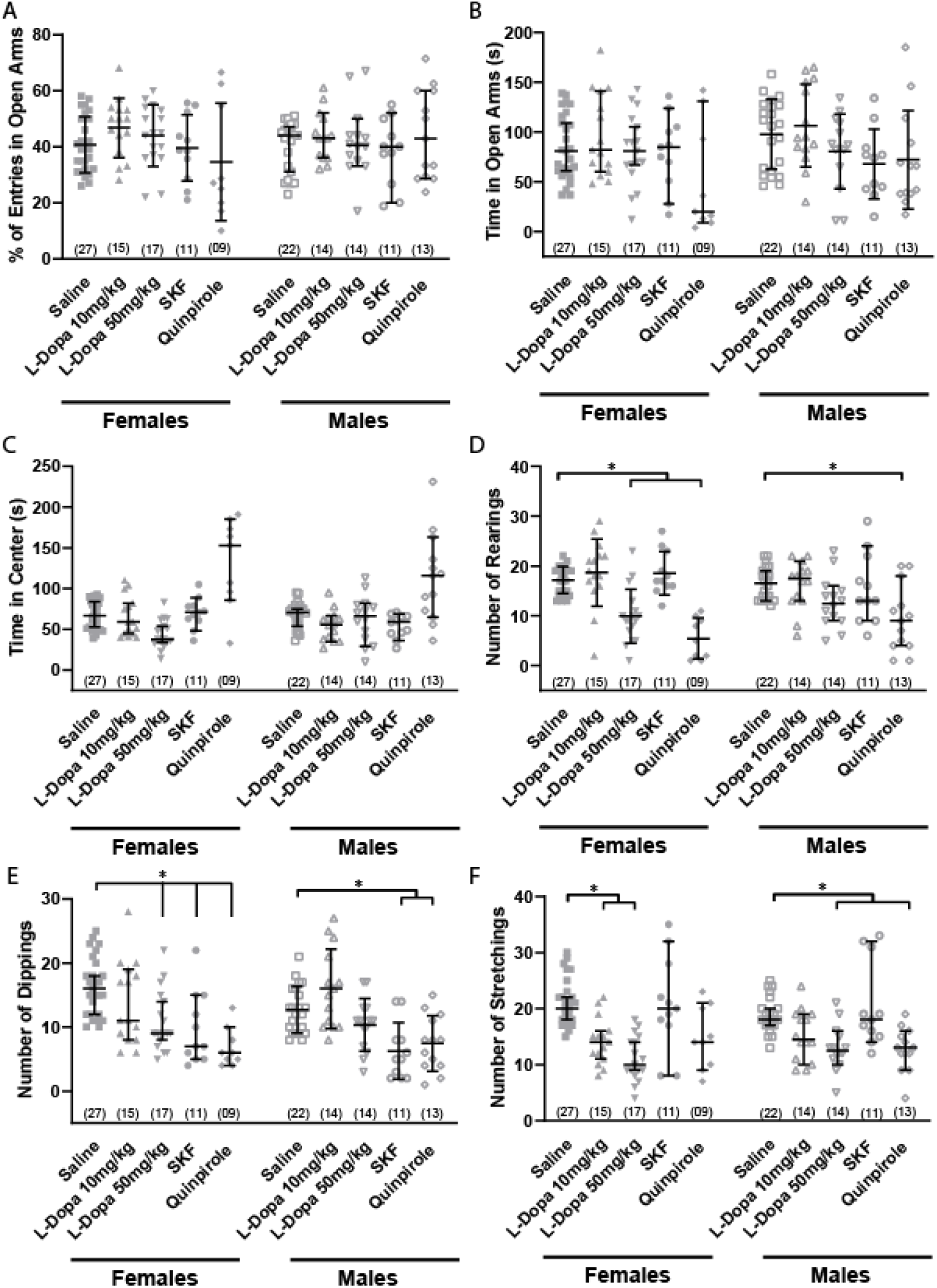
Acute increase of dopaminergic signaling during adolescence impacts exploratory behavior in female and male Swiss mice. Anxiety-like behavior of prepubertal Swiss mice (PD30) after injection of Saline, L-Dopa 10mg/kg, L-Dopa 50mg/kg, SKF-38393 or Quinpirole were tested by Elevated Plus Maze. There were no alterations in any group in (A) percentage of entries in the open arms, (B) time on open arms or (C) time in center. (D) The number of rearings was reduced in L-Dopa 50mg/kg and Quinpirole females and in Quinpirole males. (E) L-Dopa 50mg/kg, SKF and Quinpirole females, as well as SKF and Quinpirole males, showed a reduction in the number of head-dippings. (F) There was a decrease in the number of stretchings in L-Dopa 10mg/kg and L-Dopa 50mg/kg females and L-Dopa 50mg/kg and Quinpirole males. Nonparametric data are represented as median with interquartile range. Parametric data are represented as mean ± SD. For parametric data we used One Way ANOVA test and Tukey Test. For nonparametric data we used Kruskal-Wallis One Way ANOVA on Ranks and Dunn’s test. *p < 0.05.

Males from the Quinpirole administered group showed a reduction of 43% in the number of rearings (KW H=14.264, 4 d.f., Dunn’s Method, p<0.05), compared to saline males (Figure 3D). We also found a reduction of 51% in the number of dippings in SKF males (F(4,73)= 10.074, One Way Anova, Tukey, p=0.003), as well as a 41% decrease in the Quinpirole group (p=0.013) when compared to saline (Figure 3E). Additionally, we observed a decrease of 31% in the number of stretchings in L-Dopa 50mg/Kg (KW H=24.785, 4 d.f., Dunn’s Method, p<0.05) and a decrease of 33% in Quinpirole injected males (p<0.05) compared to the saline group (Figure 3F).

Afterward, we tested whether the anxiety-like behavioral consequences of acute L-Dopa and agonists administration were statistically different between sexes. We observed that the sex-related effects in the number of rearings were distinct when it comes to the Quinpirole administered group (F(4,143)= 2.159, Two Way ANOVA, Tukey, p=0.048) (Figure 4A), whereas the number of dippings was different when administered L-Dopa 10mg/kg (F(4,143)= 2.498, Two Way ANOVA, Tukey, p<0.001) (Figure 4B).

**Fig. 4.**
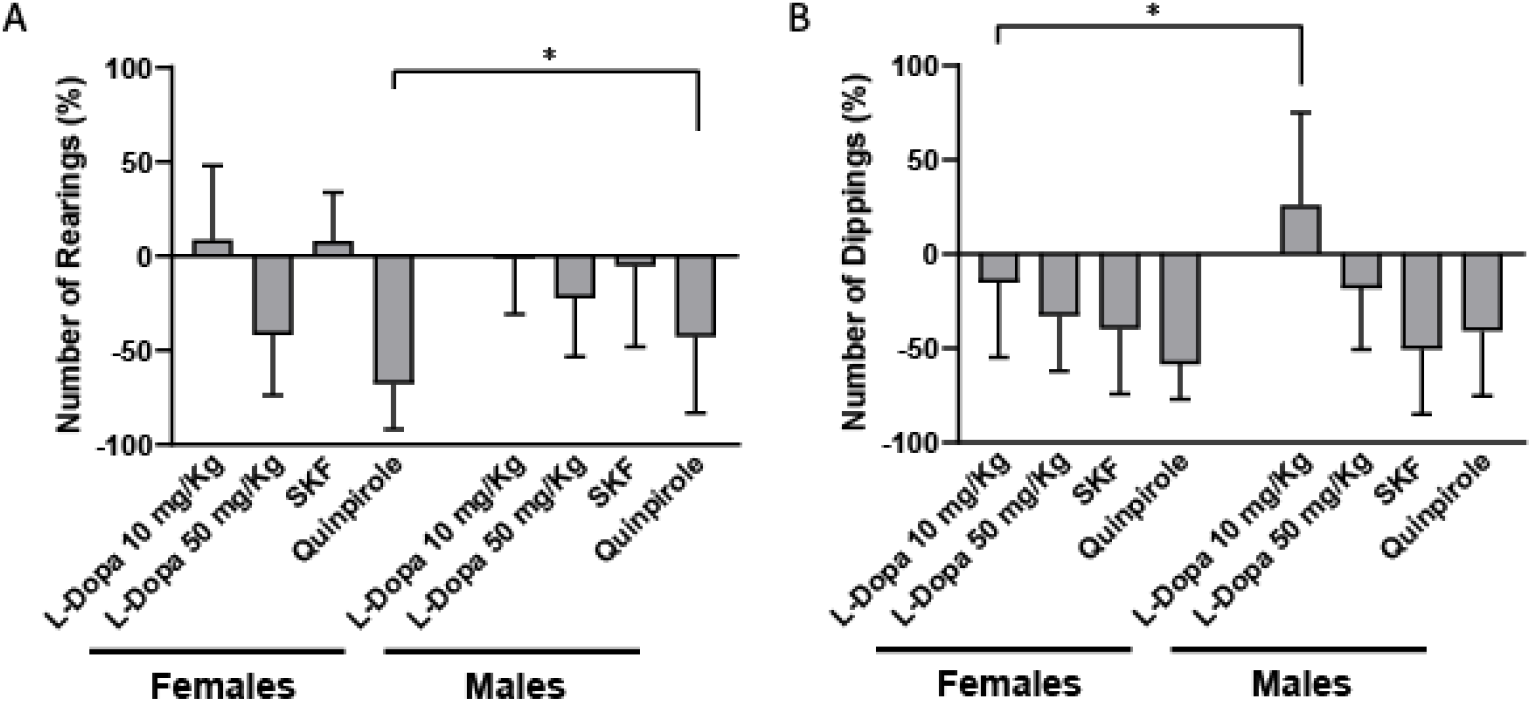
Anxiety-like and exploratory sex-specific effects following acute increase of dopaminergic signaling during pubertal phase. Differences between male and female prepubertal mice injected with Saline, L-Dopa 10mg/kg, L-Dopa 50mg/kg, SKF-38393 or Quinpirole in the Elevated Plus Maze. (A) Quinpirole females did fewer rearings than males. (B) L-Dopa 10mg/kg males had a higher number of dippings compared to females. We used the Two Way ANOVA and Tukey Test, data are represented as mean ± SD *p < 0.05.

### Increased levels of DA lead to depressive-like behavior in pubertal mice

Dopaminergic signaling is involved in the pathophysiology of depression in humans and depressive-like behavior in rodents (Dubol et al., 2020; Otte et al., 2016; Tye et al., 2013). We used FST to investigate whether the administration of L-Dopa or DA receptor D1 and D2 agonists alter depressive-like behavior in pubertal males and females. We observed a decrease of 27% in the activity time of females administered with L-Dopa 50mg/Kg (F(4,57)=3.822, One Way ANOVA, Tukey, p=0.039) compared with saline (Figure 5B). Additionally, females administered with L-Dopa 50mg/Kg showed a decrease of 34% in the immobility episodes (F(4,57)=4.054, One Way ANOVA, Tukey, p=0.024) compared with the control group (Figure 5E). Also, L-Dopa 50 mg/Kg females showed an increase of 92% in the immobility rate (KW H=18.736, 4 d.f, Dunn’s Method, p<0.05), indicating that despite fewer immobility episodes, this group stood immobile more time per episode than the saline group (Figure 5F).

**Fig. 5.**
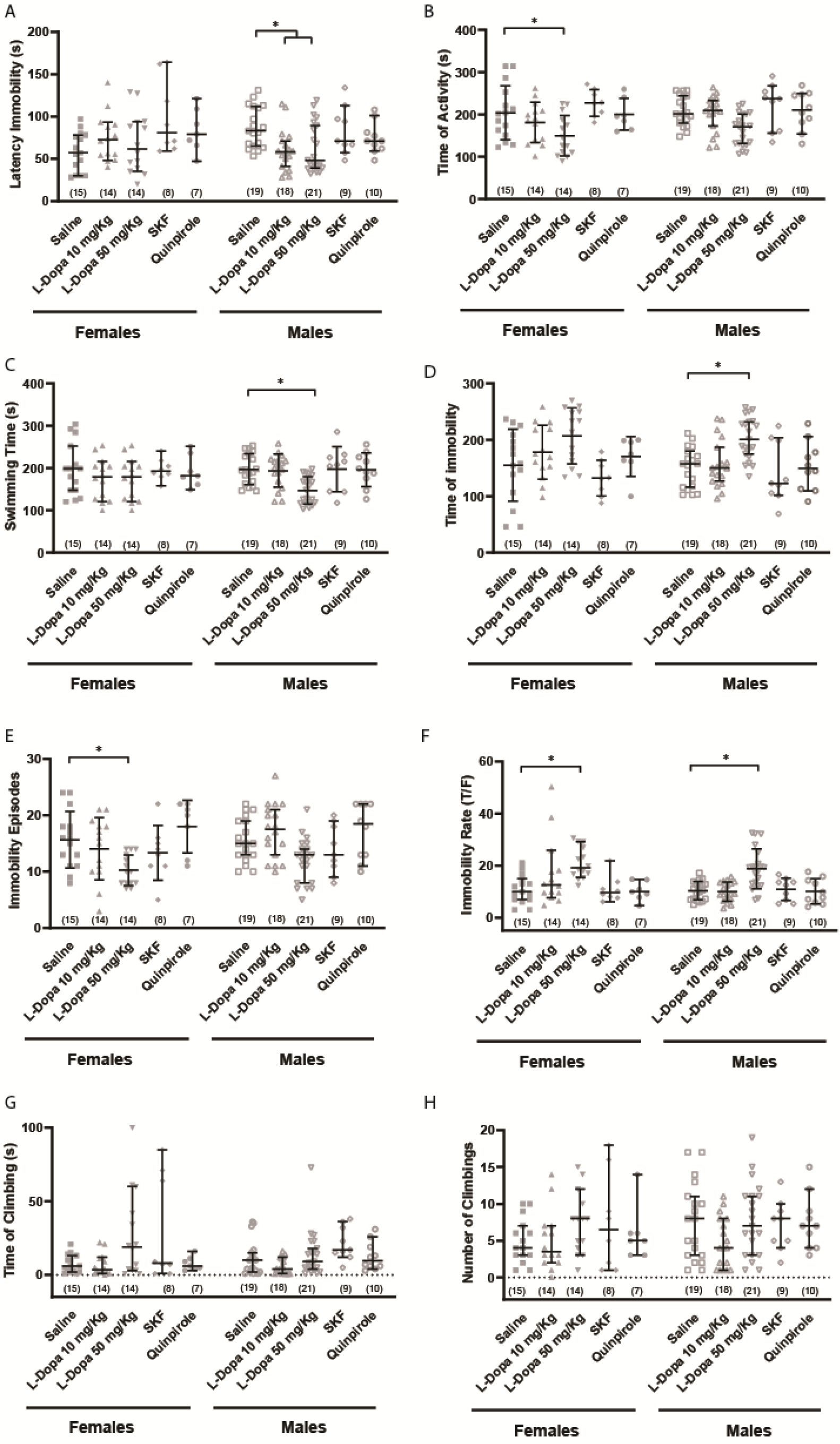
Acute increase of dopaminergic signaling during adolescence impacts depressive-like behavior in female and male Swiss mice. Depressive-like behavior of prepubertal Swiss mice (PD30) after injection of Saline, L-Dopa 10mg/kg, L-Dopa 50mg/kg, SKF-38393 or Quinpirole were tested by Forced Swimming Test. (A) L-Dopa 10mg/kg and L-Dopa 50mg/kg males decreased their latency immobility. (B) There was a decrease in time of activity of L-Dopa 50mg/kg females. L-Dopa 50mg/Kg males showed a decrease in (C) swimming time and (E) immobility episodes, as well as an increase in (D) time of immobility. (F) Both male and female L-Dopa50mg/Kg increased their immobility rate. There were no differences in (G) time of climbing nor in (H) number of climbing. Nonparametric data are represented as median with interquartile range. Parametric data are represented as mean ± SD. For parametric data we used One Way ANOVA test and Tukey Test. For nonparametric data we used Kruskal-Wallis One Way ANOVA on Ranks and Dunn’s test. *p < 0.05.

On the other hand, we observed a decrease of 31% in latency immobility in males administered with L-Dopa 10mg/Kg (KW H=14.09, 4 d.f, Dunn’s Method, p<0.05) and a decrease of 28% in L-Dopa 50mg/Kg (p<0.05) compared to the saline group (Figure 5A). We also observed a decrease of 25% in the swimming time of the L-Dopa 50mg/Kg male group (F(4,76)=6.154, One Way ANOVA, Tukey, p=0.001) (Figure 5C). Besides, we found an increase of 34% in the time of immobility of the L-Dopa 50mg/Kg males’ group (KW H= 18.125, 4 d.f, Dunn’s Method, p<0.05) compared to the saline group (Figure 5D). L-Dopa 50mg/Kg males also showed an increase of 82% in the immobility rate (F(4,76)=9.961, One Way ANOVA, Tukey, p<0.001), indicating that they spent more time immobile per episode than the saline-injected mice (Figure 5F). We found no differences in climbing time or number, neither in males and females (Figure 5G and H).

Finally, we tested the sex-related consequences of acute L-Dopa and agonists administration in the depressive-like behavior. We observed that females and males had different latency immobility in all administered groups (F(4,125)=3.379, Two Way ANOVA, Tukey, L-Dopa 10 mg/Kg p<0,001; L-Dopa 50 mg/Kg p=0,002; SKF p<0,001, quinpirole p=0,006) (Figure 6A). We also observed a difference between female and male in the immobility episodes (F(4,125)= 0.738, Two Way ANOVA, Tukey, p=0.01) and immobility rate (F(4,125)=1.435, Two Way ANOVA, Tukey, p=0.001) in L-Dopa 10mgKg groups (Figure 6B and 6C). Additionally, we found that the difference between sexes was statistically significant in the time of climbing in the L-Dopa 50mg/Kg and SKF groups (F(4,125)=2.678, Two Way ANOVA, Tukey, p<0,001; p=0.011, respectively) (Figure 6D) and in the number of climbing in L-Dopa 10 mg/Kg, L-Dopa 50 mg/Kg and SKF (F(4,125)=0.789, Two Way ANOVA, Tukey, p=0,022, p=0,002, p=0,012, respectively/ Figure 6E).

**Fig. 6.**
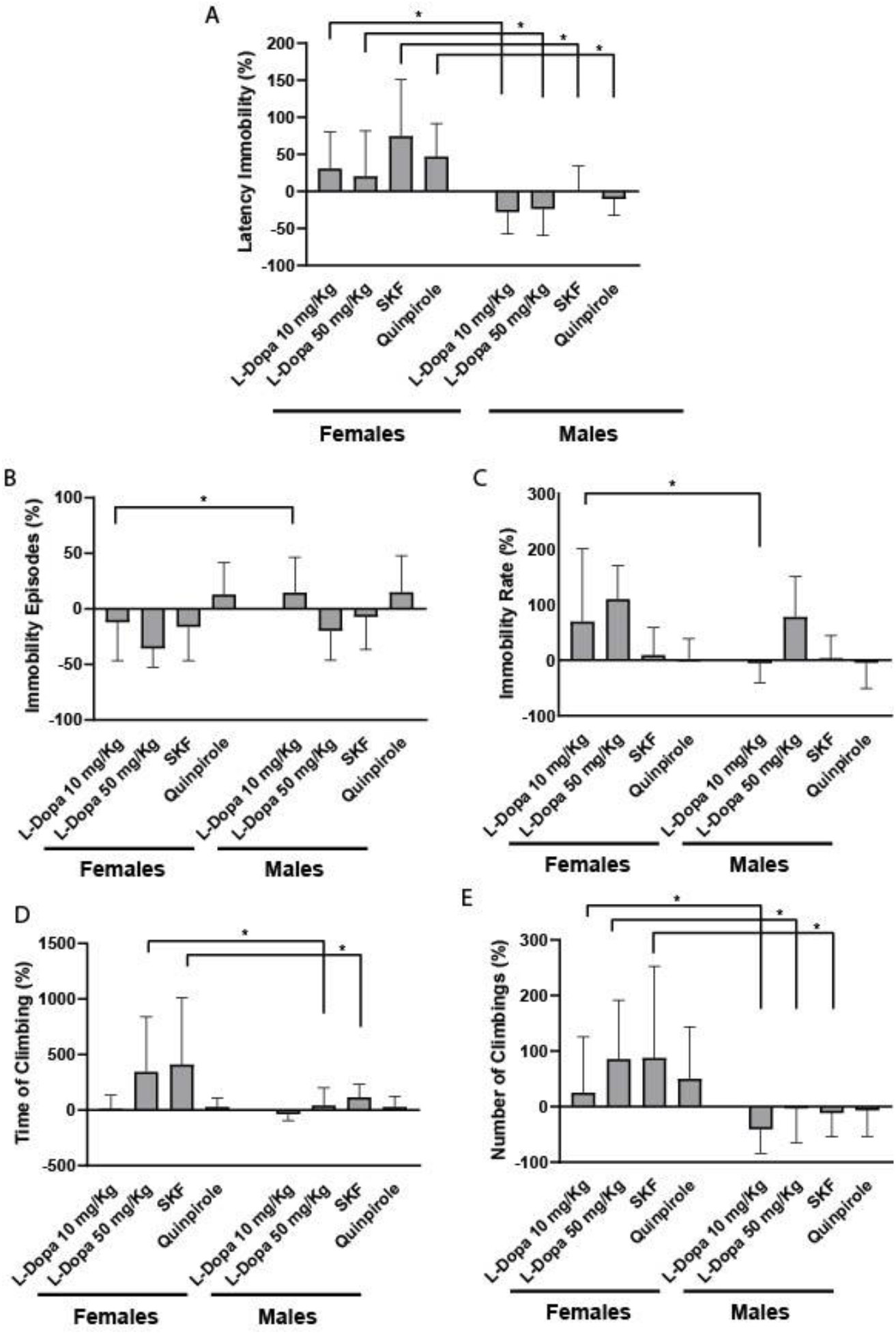
Depressive-like sex-specific effects after an acute increase of dopaminergic signaling during pubertal phase. Depressive-like differences between male and female prepubertal mice injected with Saline, L-Dopa 10mg/kg, L-Dopa 50mg/kg, SKF-38393 or Quinpirole. (A) The latency immobility was inverted in all males’ experimental groups compared to females. (B) L-Dopa 10mg/kg males presented higher numbers of immobility episodes when compared to females. (C) L-Dopa 10mg/kg males presented a lower immobility rate when compared to females. (D) L-Dopa 50mg/kg and SKF males showed decreased climbing time than males. (E) The number of climbing in L-Dopa 10mg/kg, L-Dopa 50mg/kg and SKF males were lower than in females. We used the Two Way ANOVA and Tukey Test, data are represented as mean ± SD *p < 0.05.

## Discussion

This study explores the consequences of an acute pharmacological intervention on the DAergic system in the anxiety- and depressive-like behaviors of early adolescent male and female mice. Adolescence is a developmental time window of vulnerability, with heightened sensitivity to environmental cues and an enhanced neural malleability (Brenhouse & Andersen, 2011; Casey et al., 2008; Fuhrmann et al., 2015). Interventions during this period lead to specific outcomes, not necessarily the same as those expected when studying infancy or adulthood. In this study, acute stimulation of the DAergic system was linked to decreased exploratory behavior in females, whereas males presented a depressive-like behavioral pattern, and a reduction in exploration only when exposed to an aversive context.

Scrutinizing ethological behaviors and receptor responses at the early adolescence phase is imperative. This is the last developmental window with extensive brain modifications, which can be a reason for vulnerability to environmental conditions and subsequent onset of psychiatric disorders. At this time, the brain undergoes rearrangement, maturation, and synaptic pruning (Andersen, 2003; Keshavan et al., 2014). Just as symptomatology differs between adolescents and adults, rodent behavior responses are also different. Our group’s previous work showed that early life transient increase of dopamine leads to different behavior outcomes at early adolescence and adulthood in mice. Interestingly, we evideniated a shift response at different ages (Matos et al., 2018, 2020). In accordance, other researches regarding the DAergic system describe different responses through ages (Brenhouse & Andersen, 2008; Corongiu et al., 2019; Nemoda et al., 2011; Zbukvic et al., 2017).

DAergic fluctuations during adolescence can cause behavior alterations (Padmanabhan & Luna, 2014). For example, adolescent male rats injected with Quinpirole showed compulsive and hyperactive behavior compared to control (Straathof et al., 2020); and juvenile animals show an enhancement in reinforcing effects to DAergic drugs (Adriani et al., 1998; Brenhouse & Andersen, 2008; Corongiu et al., 2019; Walker & Kuhn, 2008). The amount of evidence regarding DA signaling during adolescence is growing, counting with distinct approaches to address this particular question. DAT Knockout mice and mice lacking D2 autoreceptors have impaired mesocortical circuitry (Brami-Cherrier et al., 2014; Zhang et al., 2010) and display behavior alterations (Brami-Cherrier et al., 2014; Spielewoy et al., 2000). However, these mice lack the target protein throughout their entire life and one could not separate homeostatic effects from the consequences of a disruption of DA signaling in a specific time-window. Thus, pharmacological challenges could be a better approach to acquire the behavioral consequences of the transient increase of DA and DAergic receptor activation in adolescence.

In our study, we found a reduction in female exploratory behavior after the stimulation of DA receptors. Females injected with L-Dopa 50mg/kg decreased the total distance travelled in the Open Field Test, and L-Dopa 10, 50mg/kg and Quinpirole females also did fewer rearings, a vertical exploratory behavior (Figure 1). Although we have not found a conventional anxiety-like behavior in the Elevated Plus Maze - as there were no differences in the time spent in the open arms - our pharmacological interference in the DAergic signaling led to alterations in ethological parameters. In this test, female L-Dopa 50mg/kg and Quinpirole groups showed a reduction in vertical exploration (Figure 3). These groups plus the SKF group decreased the head-dippings, a behavior in which the animal projects its head over the edge of the open arms. Also, L-Dopa 10 and 50mg/kg females did fewer stretchings (Figure 3). The stretching can be interpreted as a risk-assessment behavior, but both the stretchings and the total number of head-dippings are also indicatives of exploratory activity (Carola et al., 2002; Rodgers et al., 1997; Rodgers & Johnson, 1995). Consistently, when we analyzed the Open Field Test by time, the decrease in the total distance travelled by female L-Dopa 50mg/kg is only present in the first 5 minutes (Figure 1). The arrival in a new environment can induce not only fear drive but also exploratory drive, especially during the first minutes, known for hyperresponsiveness (Montgomery, 1955). Adinolfi and colleagues (2018) showed that adult and adolescent DAT-KO rats process novelty differently and are less prone to explore novel environments when compared to wild type animals in a context-dependent manner (Adinolfi et al., 2018). Different behavioral aspects can influence the performance in the Open Field Test (Tanaka et al., 2012) and the exploratory activity towards novel stimuli. Therefore, the distance travelled in this specific time window, together with the decreased exploration of the Elevated Plus Maze, may indicate a diminution in the exploratory drive.

The amount of DA in the synaptic cleft and the firing of DAergic neurons may have a role in the exploratory drive. Düzel et al. (2010) proposed that the balance between tonic and phasic firing modes modulates the motivation toward novelty, considering that the tonic firing would energize exploratory behavior, leading to phasic responses after the exposure to a new environment (Düzel et al., 2010). In this work, even though there was a novel environment following DAergic stimuli, our acute interference possibly changed the interrelation of dopaminergic signaling and exploratory motivation. Although there were specific outcomes within the experimental groups, the big picture of our data shows a disruption of female exploratory-based behaviors after the pharmacological stimulation of the DAergic system.

In contrast to females, the aversive context of the Elevated Plus Maze was a determinant factor for the exploration disruption in males. Despite exploring the Open Field, males injected with Quinpirole showed fewer rearings, head-dippings, and stretchings, all ethological parameters related to exploration (Figure 3). Consistently, pharmacological stimulation of D2 receptors was associated with decreased exploratory locomotion in male adult rats (Mogenson & Wu, 1991), as well as less exploratory behavior in the Elevated Plus Maze in adult male mice (Rodgers et al., 1994). Therefore, within sex specificities, our data shows that DAergic signaling is a component of exploratory-driven behavior.

Female pubertal mice injected with L-Dopa 50 mg/Kg decreased their total time of activity in the Forced Swimming Test and spent more time immobile per episode, due to a decrease in the number of immobility episodes. However, they did not present alterations in the individual parameters of activity (swimming and climbing time) nor at immobility time (Figure 6). Although some authors did not find correlations between locomotor or anxiety-like behaviors and the Forced Swimming Test (Hilakivi & Lister, 1990; Lino-De-Oliveira et al., 2005), they usually compare classical parameters, rather than ethological ones. In contrast, some authors discuss that decreased locomotor activity can be related to false-positive results in this test (Petit-Demouliere et al., 2005; Slattery & Cryan, 2012) and stimulants like caffeine modify several parameters at the Forced Swimming Test (Costa et al., 2013). In addition, the majority of data available relies on male adult mice or rats, and may not be reproducible in females. Once we do not have enough evidence of depressive-like behavior in female mice, we believe that the alteration seen may be a consequence of the decreased exploratory behavior.

On the other hand, males injected with L-Dopa 50 mg/Kg had a decrease in latency to immobility and swimming time, as well as an increase in immobility time in the Forced Swimming Test (Figure 6). These parameters are the classic indicators of depressive-like behavior, as they are reverted by antidepressants in both rats and mice (Castagné et al., 2009; Costa et al., 2013; David et al., 2001; Porsolt et al., 1978; Porsolt et al., 1977; Slattery & Cryan, 2012). L-Dopa 50 mg/Kg males also spent more time immobile per episode compared to the control group, as a consequence of an increase in total immobility time. Thus, contrasting with females, the Forced Swimming Test suggests that early adolescent male mice presented a depressive-like behavior after administration of L-Dopa 50mg/Kg. Interestingly, specific dopamine receptor activation by SKF and Quinpirole did not alter mice behavior in this test (Figure 6). Consistently, D1, D2, and D3 agonists do not alter male mice behavior when administered alone but modulate the antidepressive responses of several Selective Serotonin Reuptake Inhibitors (Renard et al., 2001). Both haloperidol and sulpiride (respectively D1- and D2-like receptor antagonists) reversed the antidepressive effect of tramadol in the Forced Swimming Test in adult male Swiss mice (Jesse et al., 2010). The synergy of DA receptors was also demonstrated by Dwyer and collaborators (2016) in ambulatory and stereotypic behavior (Dwyer et al., 2016). These findings reflect the necessity of an integrated DAergic receptor response for behavioral outcomes.

The molecular feature of the DAergic response was not the focus of our work, but it may be responsible for such differences in early adolescence. At this time-point, DA mesocortical axons grow from the striatum to the prefrontal cortex, and this long-distance axon growth is sensitive to environmental influences (Hoops and Flores, 2017). It is also a period of increasing DA receptors in different regions (Caballero et al., 2016; Teicher et al., 1995), as well as changes in D1:D2 density ratio (Cullity et al, 2019). DAT levels follow the same pattern, but it seems to have a higher affinity uptake in early adolescence (Coulter et al., 1996; Tarazi et al., 1998; Walker & Kuhn, 2008). Thus, the interaction of genetic, epigenetic, and environmental factors at early adolescence may lead to different behavior outcomes and vulnerability to psychiatric disorders.

As well as age, sex can also influence the response to stimuli. Our study shows a sex-specific reaction to DAergic drugs: we observed a reduction of exploratory behavior in females and a depressive-like behavioral pattern in males. When compared to each other, males and females had specific responses to L-Dopa, SKF, and Quinpirole. Despite knowing that sex-associated features are probably in a spectral distribution across the population (Joel et al., 2015; Nguyen et al., 2019), understanding the physiological particularities of each sex is a major priority in terms of gathering inclusive evidences about brain development and functioning. There is an underrepresentation of female subjects in biological research, prominently in neuroscience (Beery & Zucker, 2011), possibly due to an erroneous belief that female data are more variable. Interestingly, meta-analytical studies have shown that the variability of data is not related to sex (Becker et al., 2016; Prendergast et al., 2014). Therefore, there is no scientific reason to not include female subjects in research protocols. In fact, studies show sex differences in DAergic and stress modulation of decision making (Georgiou et al., 2018), and reward systems (Becker & Chartoff, 2019), as well as in adolescence-related changes, like social behavior mediated by D1 receptors (Kopec et al., 2018), density of DAergic receptors (Cullity et al., 2019), and many others (Brenhouse & Andersen, 2011; De Bellis et al., 2001; Lenroot & Giedd, 2010). Hence, male-restricted studies are not only unsupported by evidence but also prejudicial to a proper understanding of brain function.

Many psychiatric disorders appear in late childhood, and studies show adolescence as an opportunity window for intervention and early diagnosis (Andersen, 2003; Keshavan et al., 2014; Shaw et al., 2008). Since the DAergic system is maturing at this period, it may play an important role in the pathophysiology and etiology of such disorders. Therefore, it is crucial to better understand the functionality and responses of the DAergic system to an abnormal activation in the prepubertal phase. This study highlights that sex differences in these responses appear before hormonal maturity and should be better understood to improve clinical diagnosis and treatment. Future studies are necessary in order to fully understand the ontogenetic mechanisms related to DAergic signaling in adolescence and the consequences of its disruption in such a vulnerable time-window.

## Conclusions

The acute stimulation of dopamine receptors leads to sex-specific responses even before hormonal maturity in Swiss mice: females reduced their exploratory behavior, whereas males reduced exploration only in aversive context and presented a depressive-like behavior.

## Notes

### Competing Interest Statement

The authors have declared no competing interest.

